# Gene prediction in heterogeneous cancer tissues and establishment of Least Absolute Shrinking and Selection Operator model of lung squamous cell carcinoma

**DOI:** 10.1101/737866

**Authors:** Ateeq Muhammed Khaliq, SharathChandra Rg, Meenakshi Rajamohan

## Abstract

**Background:** This study is aimed to establish a Least Absolute Shrinking and Selection Operator (LASSO) model based on tumor heterogeneity to predict the best features of LUSC in various cancer subtypes.

**Methods:** The RNASeq data of 505 LUSC cancer samples were downloaded from the TCGA database. Subsequent to the identification of differentially expressed genes (DEGs), the samples were divided into two subtypes based on the consensus clustering method. The subtypes were estimated with the abundance of immune and non-immune stromal cell populations which infiltrated tissue. LASSO model was established to predict each subtype’s best genes. Enrichment pathway analysis was then carried out. Finally, the validity of the LUSC model for identifying features was established by the survival analysis.

**Results:** 240 and 262 samples were clustered in Subtype-1 and Subtype-2 groups respectively. DEG analysis was performed on each subtype. A standard cutoff was applied and in total, 4586 genes were upregulated and 1495 were downregulated in case of subtype-1 and 5016 genes were upregulated and 3224 were downregulated in case of subtype-2. LASSO model was established to predict the best features from each subtypes, 49 and 34 most relevant genes were selected in subtype-1 and subtype-2. The abundance of tissue-infiltrates analysis distinguished the subtypes based on the expression pattern of immune infiltrates. Survival analysis showed that this model could effectively predict the best and distinct features in cancer subtypes.

**Discussion:** This study suggests that the unsupervised clustering and LASSO model-based feature selection can be effectively used to predict relevant genes which might play an important role in cancer diagnosis.

## Introduction

Lung cancer is among the most deadly cancers (Siegel, 2017). Its shows the worst survival rate when compared with colon, breast, and pancreatic cancers combined. Lung cancer is classified as non–small-cell lung cancer (NSCLC) and small-cell lung cancer (SCLC). NSCLCs are generally subcategorized into adenocarcinoma (LUAD), squamous cell carcinoma (LUSC), and large cell carcinoma. LUSC and LUAD account for 15% and 85% of all lung cancer, respectively (Inamura, 2017). Lung cancer is a highly heterogeneous disease and identification of cancer subtypes is pivotal for clinicians. Genetic mutations, cancer microenvironment, immune, and therapeutic selection pressures all dynamically contribute to tumor heterogeneity. Heterogeneity may lead to cells with a differential molecular signature within single tumor tissue and in some cases, it may contribute to therapy resistance (Beca F, 2016; Bolck et al., 2019). Therefore, deciphering LUSC cancer heterogeneity will have a major impact in designing precision medicine strategy. Heterogeneous data suffers from a large number of covariates, and identification of variable selection is necessary to obtain more accurate predictions with a large number of covariates.

Over the past decades, many computer-aided diagnostic models have been used for predicting the risk of a variety of cancers, such as logistic regression, Cox proportional hazard model, Artificial neural networks, decision trees and Support vector machines (Bartfay, Mackillop & Pater, 2006; Ayer et al., 2010; Jiang & Ching, 2012; ZHU et al., 2013).Previous studies indicate standard stepwise selection approaches which are not best for regression models with a very large number of covariates (Houssami et al., 2004). Alternatively, least absolute shrinkage and selection operator (LASSO), has received much attention for identification and selection of best variables. LASSO was first formulated by Robert Tibshirani in 1996 (Tibshirani, 1996). It is a powerful method that performs two main tasks: regularization and feature selection. LASSO estimates the regression coefficients by maximizing the log-likelihood function with the constraint that the sum of the absolute values of the regression coefficients, ∑j=1kβj, is less than or equal to a positive constant *s*.

In this study, we downloaded the RNASeq data for LUSC cancer samples from The Cancer Genome Atlas (TCGA) database. We differentiated the samples based on clusters into two subtypes to study the tumor heterogeneity. Differentially expressed genes (DEGs) were identified between two subtypes and normal groups, followed by predicting relevant variables that are associated with the response variable using the LASSO model and validating the variables using survival analysis. We estimated the population abundance of tissue-infiltrating immune and stromal cell populations in each subtype to decipher the inflammatory, antigenic, and desmoplastic reactions occurring in cancer tissue. Our study provides new insight into tumor heterogeneity and its importance in sample classification for predicting of biomarkers of LUSC cancer.

## Materials & Methods

### Data source

The RNASeq data of Lung Squamous cell cancer, including 505 LUSC samples, and 49 normal samples were downloaded from the TCGA database (https://portal.gdc.cancer.gov/) in May 2019. All the raw, preprocessed data and supporting files can be accessed at https://bitbucket.org/lusc_data/supporting_data/src/master/.

### Data preprocessing and grouping

Based on the clinical data, the LUSC cancer samples downloaded from TCGA database were divided into two sets, the first set was divided into 114 low-risk samples and 390 high-risk sample groups according to the AJCC Cancer Staging (https://cancerstaging.org/). The second set (set2) was divided into 505 Primary solid Tumor samples and 49 Solid Tissue Normal samples. We calculated a variance stabilizing transformation (VST) from the data and transformed the counts yielding a matrix of values approximately homoskedastic.

### Molecular subtyping analysis

Feature dimension reduction was needed to remove irrelevant features and to reduce noises, we used median absolute deviation (MAD) method and the features with MAD>0.5 were selected from set 2 groups. Consensus clustering (CC) (Monti et al., 2003) was used for the identification of subtypes on set 2 group. Silhouette width (Rousseeuw, 1987) was used to validate sample clustering to its identified subtype compared to other subtypes.

### Differential gene expression analysis

Differential gene expression was assessed by using the DESeq2 package (Love MI, Huber W, 2014) (Version 1.24.0, https://bioconductor.org/packages/release/bioc/html/DESeq2.html) on set1 (High-Risk samples Vs. Low-risk samples) and set2 (Subtype-1 vs. Normal and Subtype-2 vs. Normal). Log2 fold change > 2 and P-value <0.05 were used as the cut-off values to identify the DEGs.

### Construction of the LASSO Model

Glmnet Package (Friedman J, Hastie T, 2010) (Version 2.0-18, https://cran.r-project.org/web/packages/glmnet/index.html) was used to fit a generalized linear model via penalized maximum likelihood, LASSO model was established (Least Absolute Shrinkage and Selection Operator) on the DEGs from individual Subtype-1 and Subtype-2 cancer samples. We built a single pass (single fold) lasso-penalized model and performed 10-fold cross-validation to identify the best predictor.

### Survival Analysis

To find clinically or biologically meaningful biomarkers Kaplan-Meier survival curves (E. L. Kaplan & Paul Meier, 1958) were generated by selecting the best predictors from individual subtypes. Kaplan-Meier curves were generated using the TRGAted (Nicholas Borcherding, Nicholas L. Bormann, Andrew P. Voigt, 2018)(https://github.com/ncborcherding/TRGAted) package implemented in R.

### Quantification of the absolute abundance of eight immune and two stromal cell populations

We estimated the abundance of tissue-infiltrating immune and non-immune stromal cell populations in Subtype-1 and Subtype-2 samples. MCP-counter (Etienne Becht, Nicolas A. Giraldo, Laetitia Lacroix, Bénédicte Buttard, Nabila Elarouci, Florent Petitprez, Janick Selves, Pierre Laurent-Puig, Catherine Sautès-Fridman, 2016) (https://github.com/ebecht/MCPcounter) Package was used to estimate the Microenvironment Cell Populations. VST normalized gene expression matrix was used for the estimation of an immune and stromal cell population.

### Gene classification and enrichment analyses

clusterProfiler (Yu G, Wang L, Han Y, 2012) (Version 3.12.0, http://bioconductor.org/packages/release/bioc/html/clusterProfiler.html) was used to annotate the DEGs from Subtype-1 and Subtype-2 groups to biological processes, molecular functions, and cellular components in a directed acyclic graph structure with a q-value cutoff of 0.2, Kyoto Encyclopedia of Genes and Genomes (KEGG) (Kanehisa M, Goto S, Furumichi M, Tanabe M, 2010) was utilized to annotate genes to pathways, and Disease Ontologies.

## Results

### Identification of DEGs in High-risk LUSC tumors

The genes with p-value cutoff < 0.05 and log2 fold change > 2 were considered to be differentially expressed. A total of 22 genes were differentially expressed between high risk and low-risk samples, which includes 4 downregulated and 18 upregulated genes. *Figure 1* displays the heat map of the risk-related DEGs. Which is suggestive of similar gene expression pattern in both groups, which makes it difficult to classify the samples on the gene expression pattern. PC analysis shows the homogeneity of the data between the High and Low-risk group (*Fig 2*).

**Figure 1:**
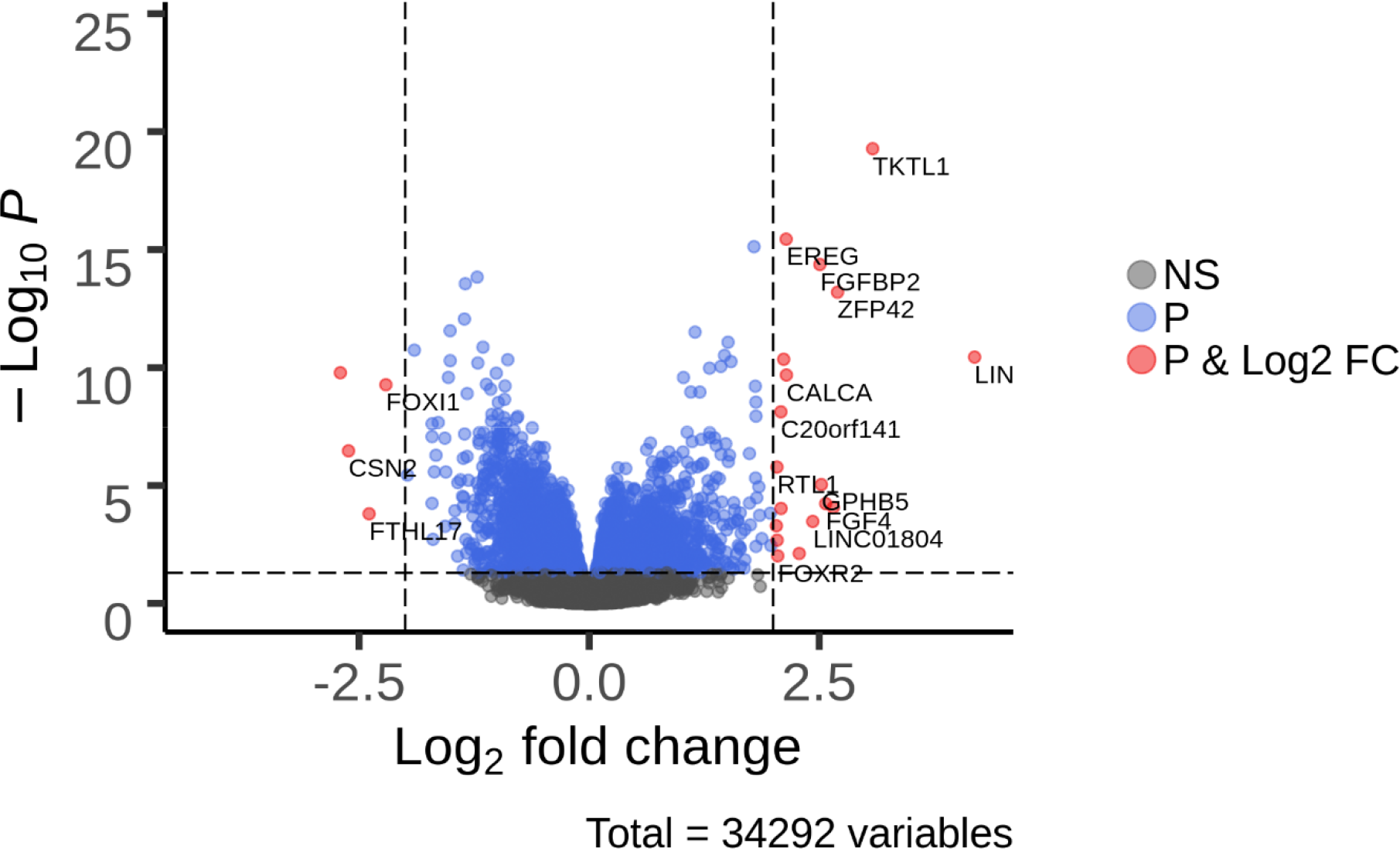
Volcano plot of differentially expressed genes (DEGs) in High risk Vs. low risk samples

**Figure 2:**
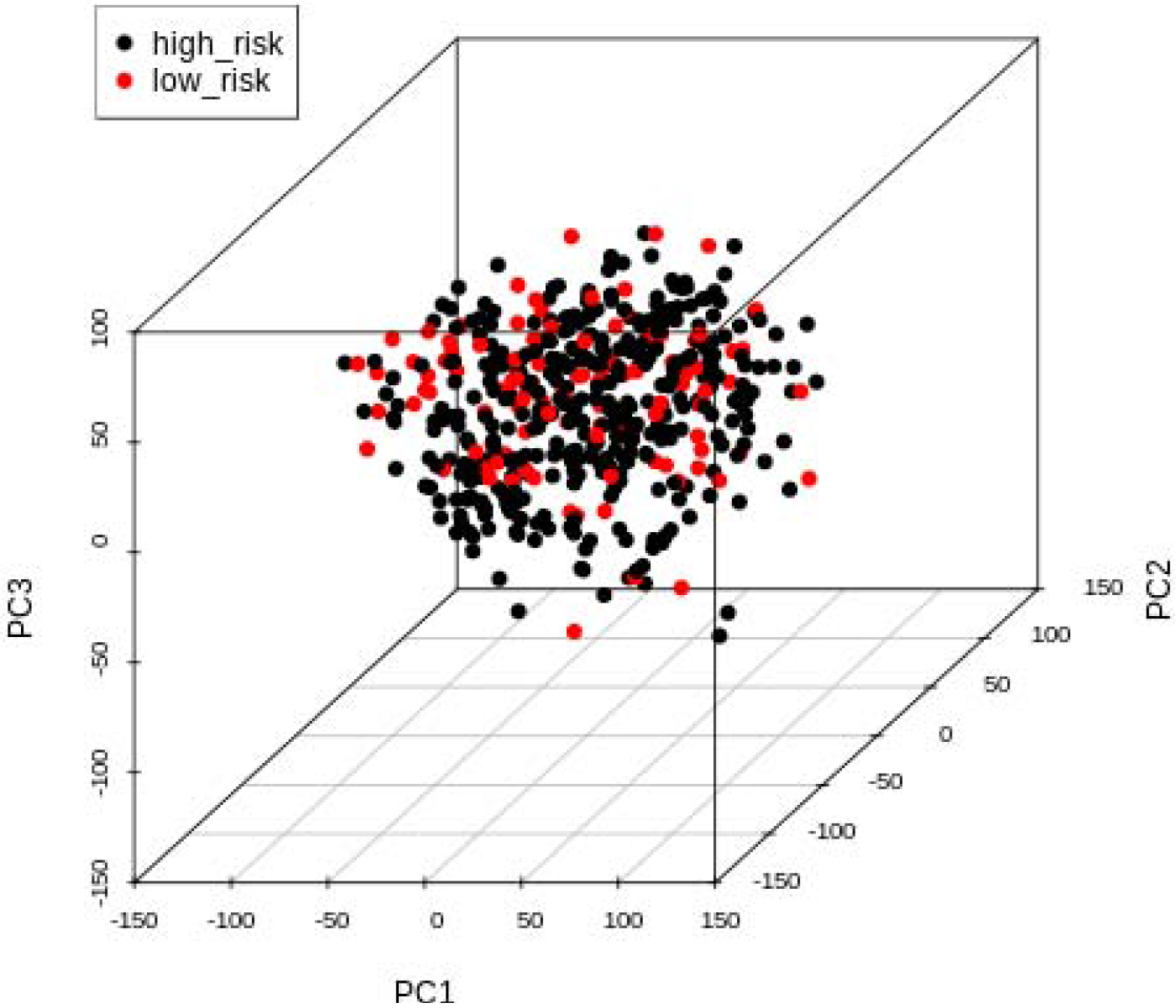
Principal component analysis for High risk and Low risk samples

### Molecular cancer subtype identification in LUSC Tumor samples and validation of clusters

We used an unsupervised clustering method Consensus clustering (CC) (Monti et al., 2003). CC method is most widely used for subtype discovery in high dimensional datasets. We used settings of the agglomerative hierarchical clustering algorithm using Pearson correlation distance. Two distinct clusters were discovered in our datasets, 240 and 262 samples were clustered in Subtype-1 and Subtype-2 groups respectively. (*Table S1 and S2*) We have validated consistency within clusters of data using Silhouette Plot (Rousseeuw, 1987). The Average Silhouette width for our generated clusters is 0.68 (*Fig 3, 4 and 5*).

**Figure 3:**
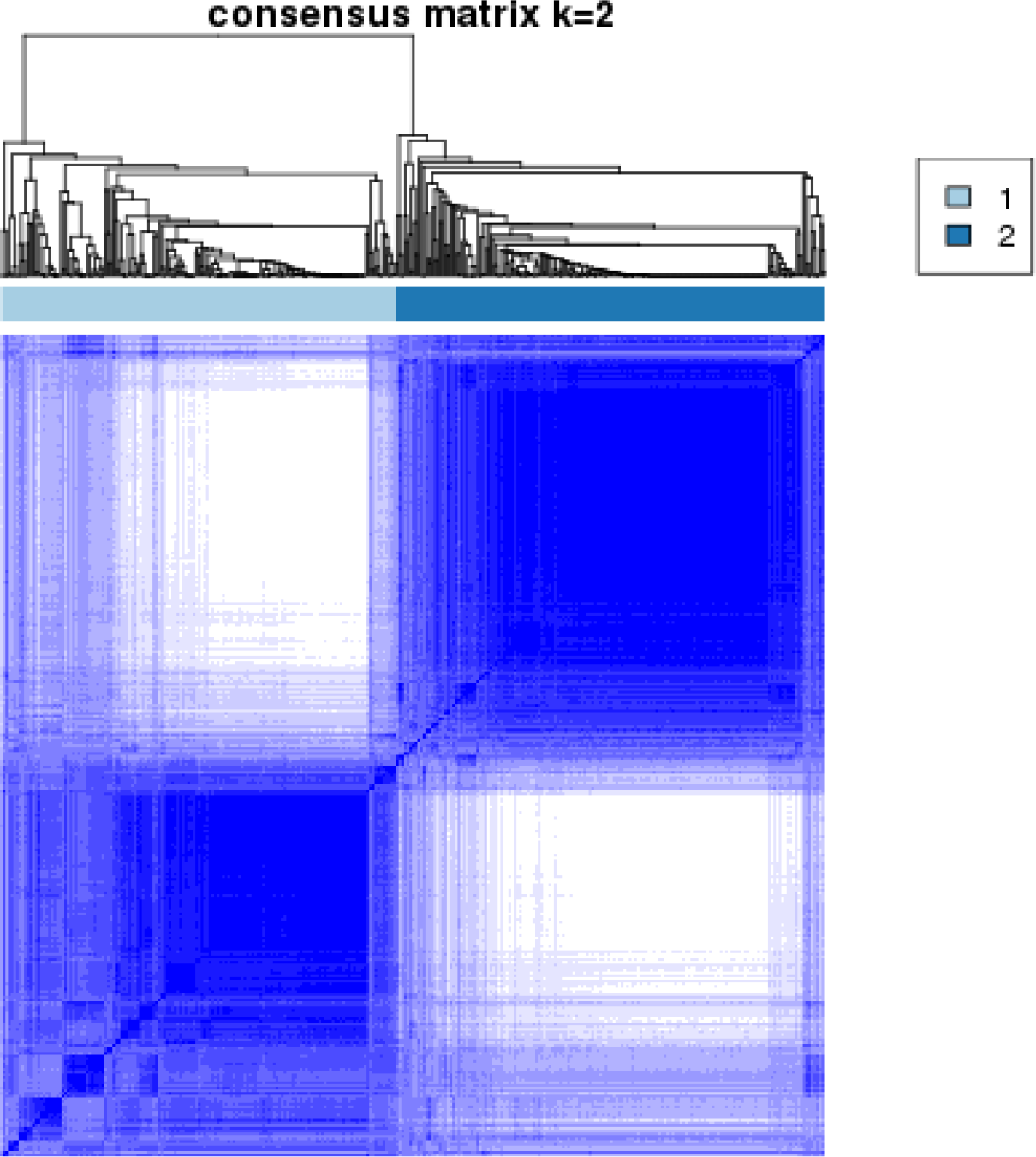
Consensus clustering results for LUSC samples

**Figure 4:**
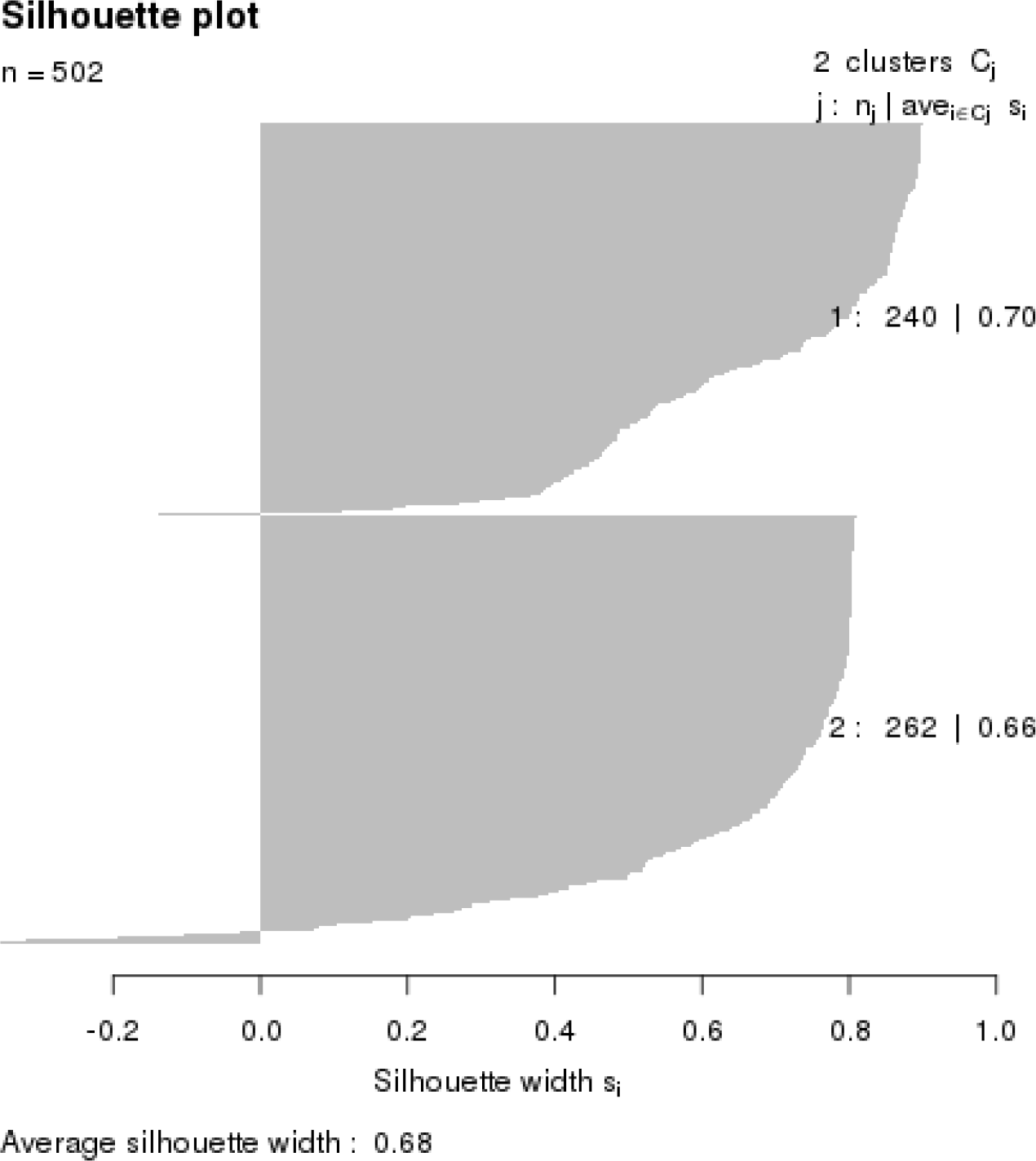
Silhouette width for Cancer subtype Validation

**Figure 5:**
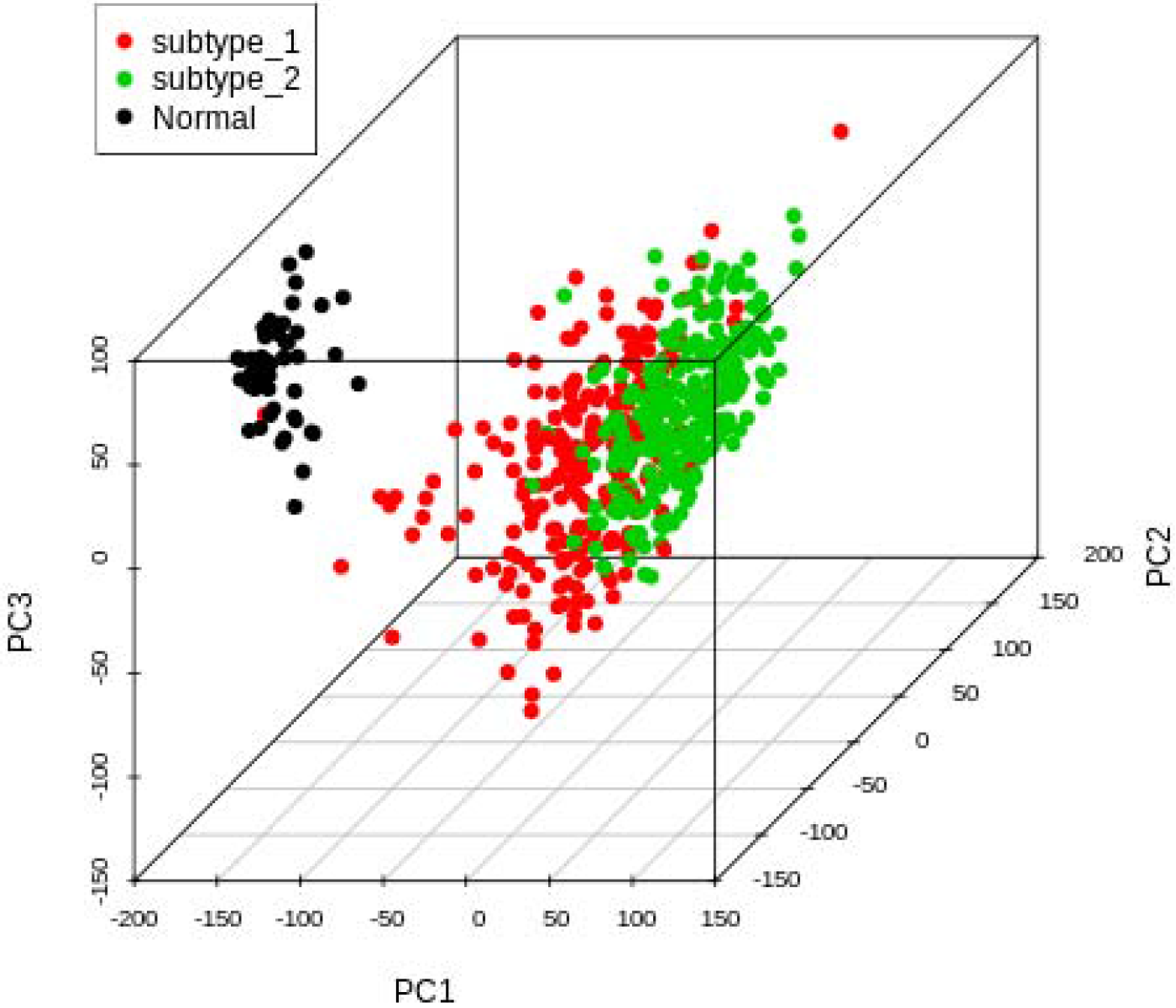
Principal component analysis for LUSC samples

### Identification of DEGs in Subtype-1 and Subtype-2 LUSC Tumors

We compared the subtype-1 and subtype-2 with the normal samples and based on the p-value cutoff < 0.05 and log2 fold change > 2 we identified significant DEGs. 4586 genes were upregulated and 1495 were downregulated in case of subtype-1 (*Fig 6*) and 5016 genes were upregulated and 3224 were downregulated in case of subtype-2 (*Fig 7*) shows differential expression pattern in subtype-1 and subtype-2. The DEGs in both subtypes were used for building LASSO predictive model and for the identification of best predictor genes.

**Figure 6:**
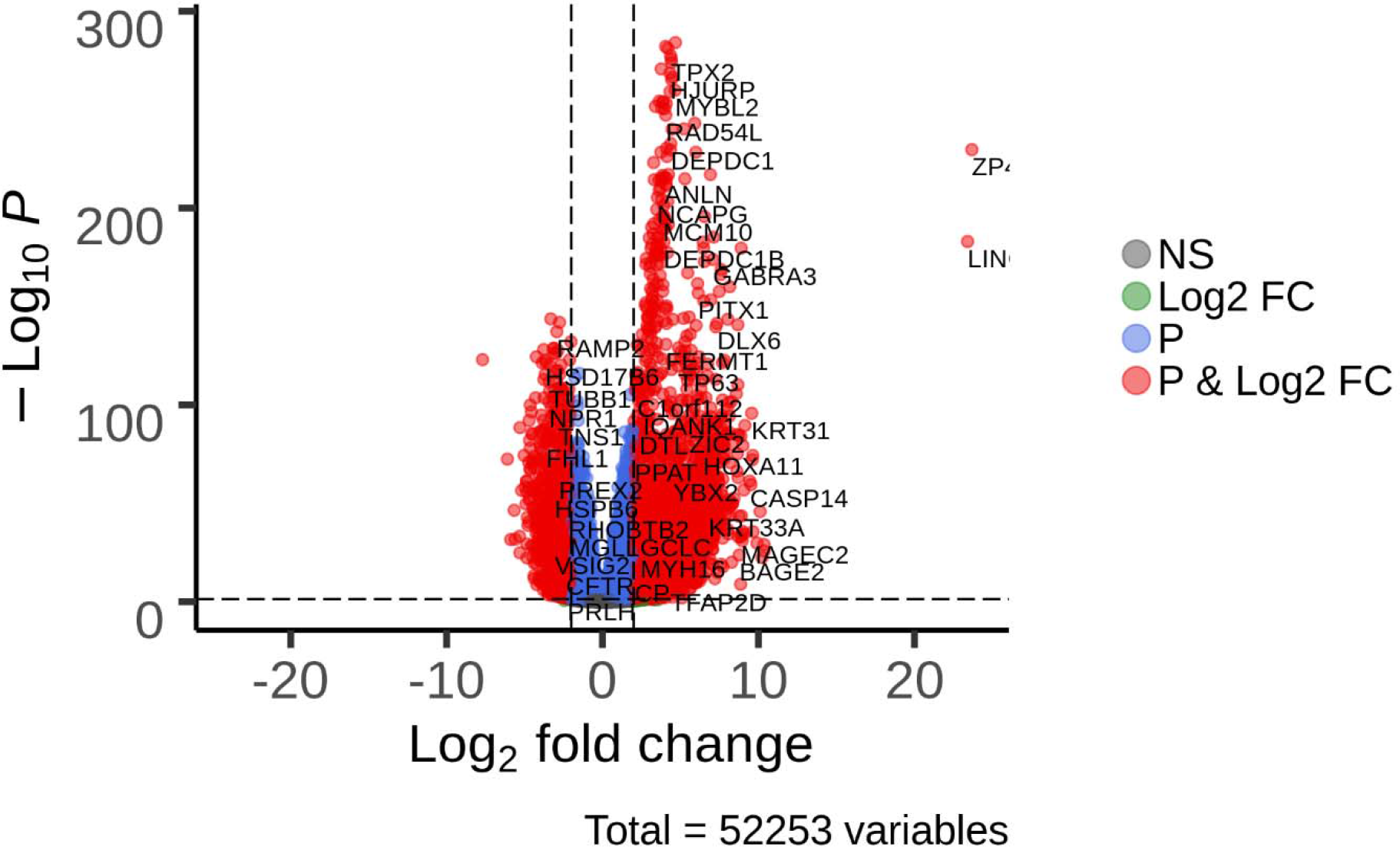
Differential gene expression in subtype 1 Vs. Normal samples

**Figure 7:**
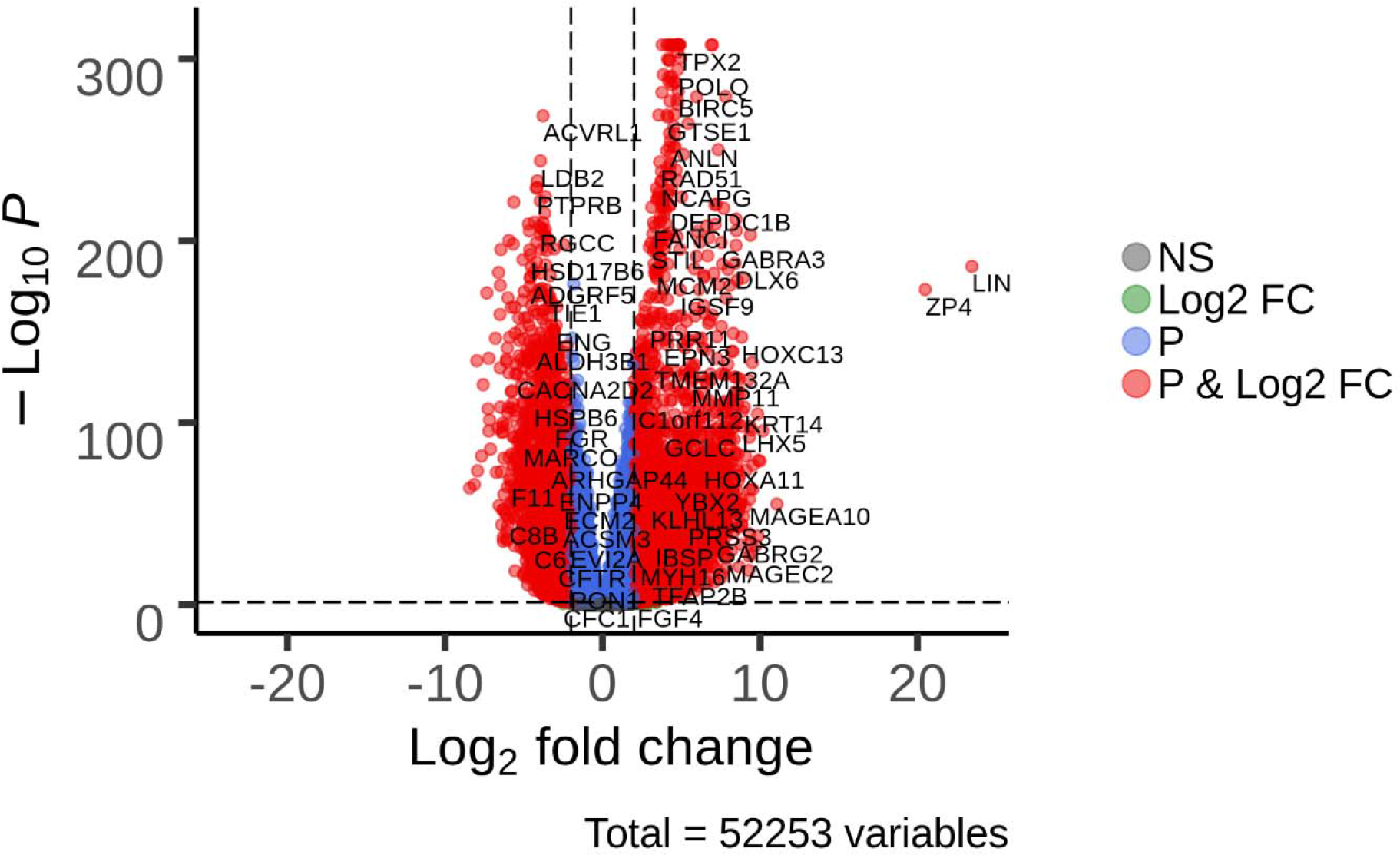
Differential gene expression in subtype 2 Vs. Normal samples

### LASSO model for identification of best predictive genes

LASSO (Least Absolute Shrinkage and Selection Operator) was first formulated by Robert Tibshirani in 1996 (Tibshirani, 1996). It is a powerful method that performs two main tasks: regularization and feature selection. RNASeq datasets are high dimensional datasets, with smaller sample size and a large number of features also called small-n-large-p datasets (p >> n). High dimensional data will be sparse and only a few features affect the response variable and LASSO is known to identify the best features that affect the response variable. We deal with a p >> n situation for feature selection in our Subtype-1 and Subtype-2 datasets, thus probably not all DEGs are relevant for the identification of features which affect the response variable. The purpose of our analysis is to identify the feature selection task and underline which genes are more relevant to predict and to classify them as biomarkers, to do so we have used the LASSO method.

The result shows the trends of the 49 and 34 most relevant features selected by our model in subtype-1 and subtype-2 LUSC cancer respectively *(Fig S1 and S2)*. The next step would be to find the most appropriate values for λ for our LASSO model. We analyzed the λ value using the 10 fold cross-validation *(Fig S3 and S4),* between λ min that gives minimum mean cross-validated error or λ1se that gives a model within one standard error of the minimum. Using this analysis we obtained the most relevant genes which are unique to subtype-1 and subtype-2 in the detection of a LUSC cancer. A list of best-predicted genes available for each cancer subtypes is shown in *Table 1* and *Table 2*.

**Table 1:** Most relevant genes in established by LASSO model in Subtype-1 of LUSC

| **Genes** | **coefficients** |
| --- | --- |
| GAL | 0.003457 |
| MYCT1 | 0.003006 |
| IMPDH1P8 | 0.001834 |
| TCF21 | 0.001811 |
| FOXA2 | 0.001753 |
| ROR1 | 0.001641 |
| NR3C2 | 0.001566 |
| GPD1 | 0.001236 |
| PPIAP45 | 0.001113 |
| LINC01977 | 0.000995 |
| GPR19 | 0.000891 |
| RTBDN | 0.000838 |
| PGM5 | 0.000726 |
| AFF3 | 0.000712 |
| LINC01572 | 0.000662 |
| TMEM249 | 0.000385 |
| TNNC1 | 0.000216 |
| CAGE1 | 0.000165 |
| TFAP2A | 2.63E-05 |
| PGM5 | -2.29E-05 |
| HRCT1 | -3.03E-05 |
| C1orf87 | -8.42E-05 |
| DPYSL2 | -0.00011 |
| NEK5 | -0.00014 |
| NKAPL | -0.00016 |
| RBMY1KP | -0.00017 |
| TGFBR2 | -0.00024 |
| EIF4EBP1 | -0.00051 |
| CENPF | -0.0006 |
| RPL31P40 | -0.0006 |
| FFAR4 | -0.00061 |
| LINC01863 | -0.0007 |
| GAS2L2 | -0.0007 |
| KCNA4 | -0.00079 |
| DUSP27 | -0.00079 |
| LINC00670 | -0.0009 |
| CAV3 | -0.00101 |
| MIR4717 | -0.00114 |
| PECAM1 | -0.00139 |
| LINC00891 | -0.00154 |
| HBM | -0.00157 |
| GP9 | -0.0016 |
| LINC02016 | -0.00169 |
| HELT | -0.0017 |
| OR6N1 | -0.0021 |
| OR6K4P | -0.00346 |
| CELF2 | -0.0037 |
| LINC02058 | -0.00486 |
| LINC00710 | -0.00499 |

**Table 2:** Most relevant genes in established by LASSO model in Subtype-2 of LUSC

| Genes | Coefficient |
| --- | --- |
| MIR3131 | -0.01287 |
| LINC00844 | -0.01122 |
| RXFP2 | -0.00708 |
| PDZRN3.AS1 | -0.00704 |
| GYPB | -0.00534 |
| KHDRBS2.OT | -0.00515 |
| DPPA3P2 | -0.00496 |
| HBM | -0.00425 |
| RPL31P40 | -0.00424 |
| LINC00710 | -0.00408 |
| SYNE1.AS1 | -0.00382 |
| GP9 | -0.00296 |
| PGM5.AS1 | -0.00277 |
| MIR6071 | -0.00207 |
| LINC01070 | -0.00201 |
| LINC01985 | -0.00184 |
| LINC02435 | -0.00152 |
| KCNA10 | -0.00119 |
| OR6K3 | -0.00113 |
| MIR4717 | -0.00094 |
| LINC02016 | -0.00077 |
| LINC00670 | -0.00066 |
| NCAPGP2 | -0.00063 |
| GGTLC3 | -0.00055 |
| GUCA2A | -0.00038 |
| ART1 | -0.0003 |
| ACSM2B | -0.00021 |
| GPIHBP1 | 2.05E-05 |
| ATOH8 | 8.75E-05 |
| TCF21 | 0.000133 |
| ADGRD1 | 0.000279 |
| MAMDC2 | 0.000348 |
| ABCA8 | 0.000531 |
| SFTA1P | 0.001316 |

### Analysis of the microenvironment of Subtype-1 and Subtype-2 LUSC cancer

The abundance of tissue-infiltrating immune and non-immune stromal cell populations is highly informative. It has been shown that the extent of infiltrating immune cells is associated with disease prognosis (Becht et al., 2016). T-cell infiltrates, endothelial cells and fibroblasts are associated with a favorable outcome and also poor prognosis in some cancer types (Pagès F, Berger A, Camus M, Sanchez-Cabo F, Costes A, Molidor R, Mlecnik B, Kirilovsky A, Nilsson M, Damotte D, Meatchi T, Bruneval P, Cugnenc P, Trajanoski Z, Fridman W, 2005; Galon J, Costes A, Sanchez-Cabo F, Kirilovsky A, Mlecnik B, Lagorce-Pagès C, Tosolini M, Camus M, Berger A, Wind P, Zinzindohoué F, Bruneval P, Cugnenc P, Trajanoski Z, Fridman W, 2006; Fridman WH, Pagès F, Sautès-Fridman C, 2012; Giraldo NA, Becht E, Pagès F, Skliris G, Verkarre V, Vano Y, Mejean A, Saint-Aubert N, Lacroix L, Natario I, Lupo A, Alifano M, Damotte D, Cazes A, Triebel F, Freeman GJ, Dieu-Nosjean M, Oudard S, Fridman WH, 2015). To understand the immunological microenvironment in our expression subset-1 and subset-2 we used MCP-counter method as described by Becht et al (Etienne Becht, Nicolas A. Giraldo, Laetitia Lacroix, Bénédicte Buttard, Nabila Elarouci, Florent Petitprez, Janick Selves, Pierre Laurent-Puig, Catherine Sautès-Fridman, 2016). The estimations consist of single sample scores which are computed on each sample independently in two subtypes. The heatmap shown in *Figure 8* clearly distinguish our subtype-1 and subtype-2 into two different categories based on tissue-infiltrating immune and non-immune stromal cell populations. Subtype-1 shows clear increase in CD8 T cells, Cytotoxic lymphocytes and Natural killer cells and Subtype-2 shows decreased levels of T-cells, macrophages, B cells, and natural killer (NK) cells, as well as endothelial cells and fibroblasts. Our study clearly distinguishes LUSC subtypes based on their inflammatory and stromal profiles and Subtype-1 LUSC samples show increased expression of immunological markers than Subtype 2 LUSC samples.

**Figure 8:**
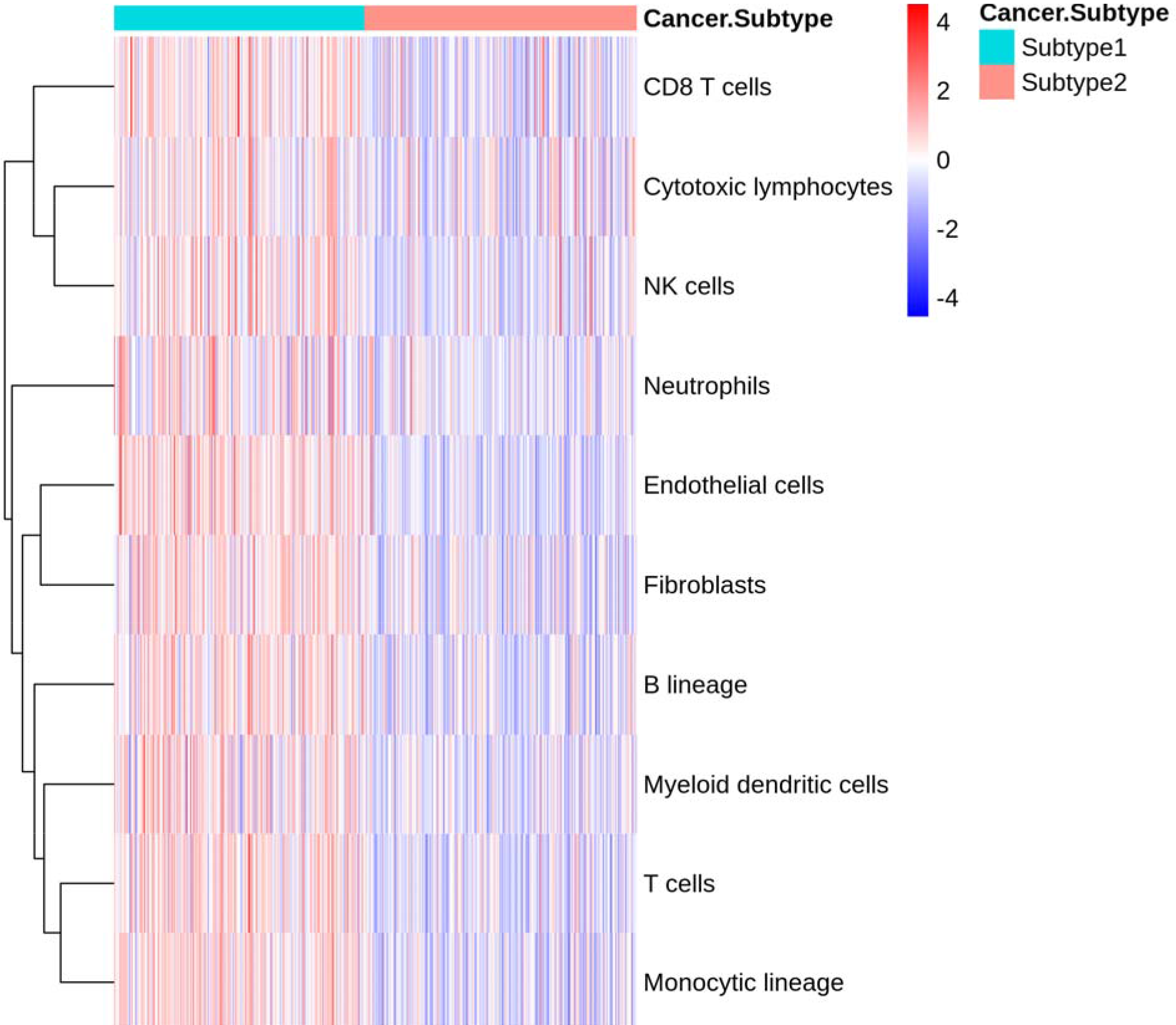
Estimating the population abundance of tissue-infiltrating immune and stromal cell populations using gene expression in Subtype-1 and Subtype-2 samples

### Disease pathway analysis

KEGG pathway analysis for Subtype-1 and Subtype-2 revealed many significant cancer pathways, including genes involved in Focal adhesion, mTOR signaling pathway, Axon guidance pathway, Cellular senescence pathway, ErbB signaling pathway, Longevity regulating pathway, HIF-1 signaling pathway, AMPK signaling pathway, Pancreatic cancer, Chronic myeloid leukemia, Colorectal cancer, TGFbeta signaling pathway etc. (*Table 3 and Table 4*). Genes such as EIF4EBP1, FOXA2, PECAM1, TGFBR2, TNNC1, ACSM2B and ABCA8 picked up by our model plays a pivotal role in cancer regulatory pathways (*Fig 9 and 10*). Both Subtype-1 and Subtype-2 groups showed distinct biological processes and cellular components after GO Enrichment Analysis of the gene sets from Subtype-1 and Subtype-2 groups. *(Fig. S5, S6, S7 and S8 and Table S3, S4, S5 and S6)*.

**Table 3:**
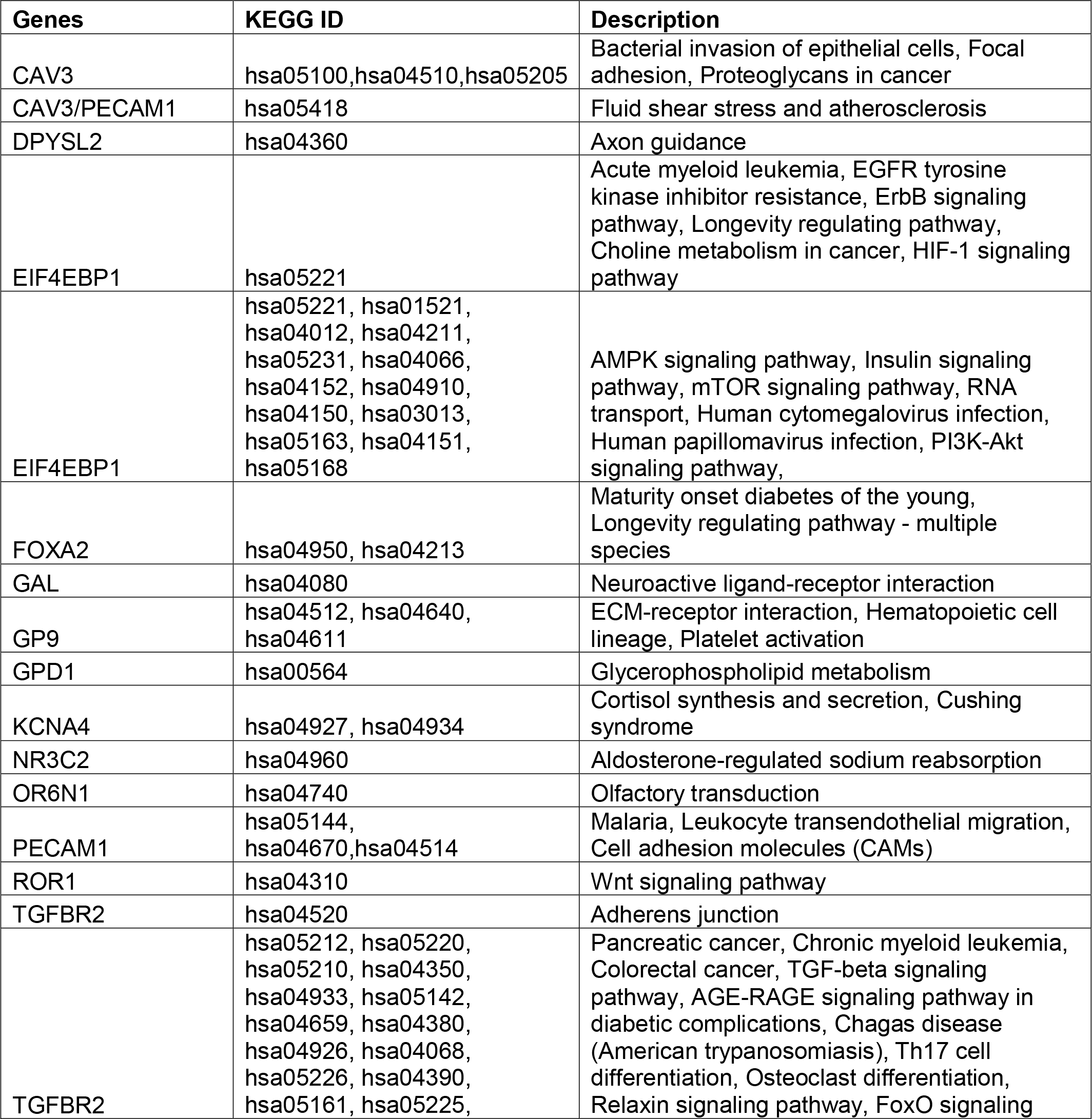
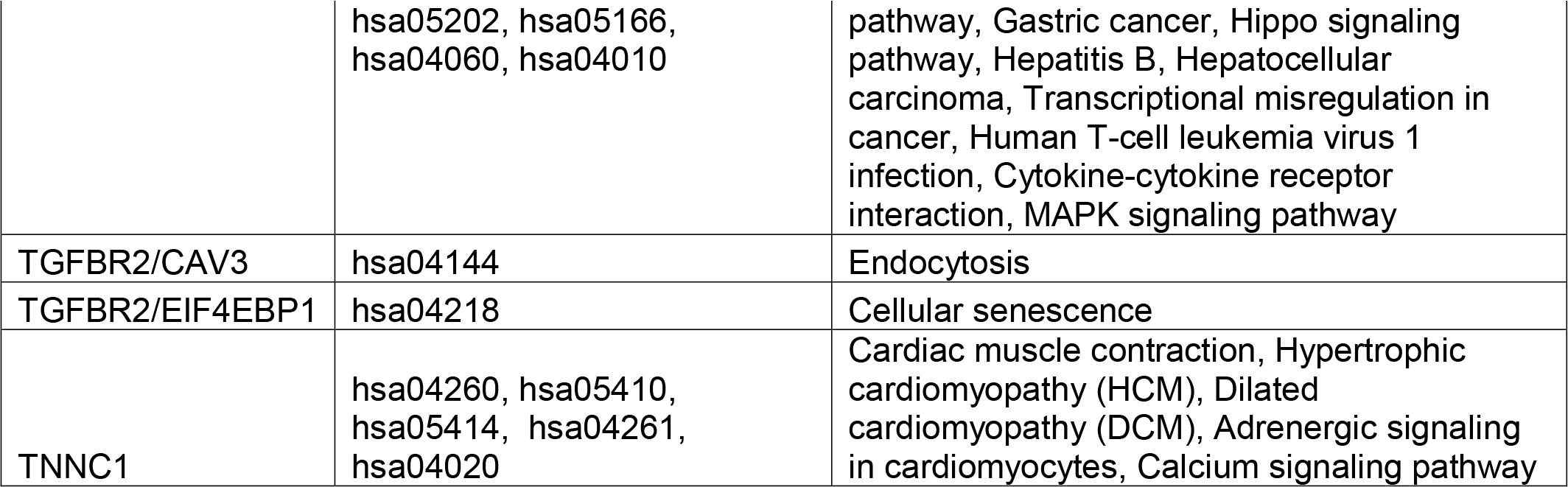
KEGG pathway analysis for Subtype-1 group

**Table 4:** KEGG pathway analysis for Subtype-2 group

| Genes | ID | Description | pvalue | p.adjust | qvalue |
| --- | --- | --- | --- | --- | --- |
| ACSM2B | hsa00650 | Butanoate metabolism | 0.021127 | 0.110183 | 0.077321 |
| ABCA8 | hsa02010 | ABC transporters | 0.033772 | 0.110183 | 0.077321 |
| GYPB | hsa05144 | Malaria | 0.036728 | 0.110183 | 0.077321 |
| GP9 | hsa04512 | ECM-receptor interaction | 0.06515 | 0.121208 | 0.085058 |
| GP9 | hsa04640 | Hematopoietic cell lineage | 0.071609 | 0.121208 | 0.085058 |
| GP9 | hsa04611 | Platelet activation | 0.090762 | 0.121208 | 0.085058 |
| RXFP2 | hsa04926 | Relaxin signaling pathway | 0.094273 | 0.121208 | 0.085058 |
| RXFP2 | hsa04080 | Neuroactive ligand-receptor interaction | 0.231252 | 0.260158 | 0.182567 |
| OR6K3 | hsa04740 | Olfactory transduction | 0.296107 | 0.296107 | 0.207794 |

**Figure 9:**
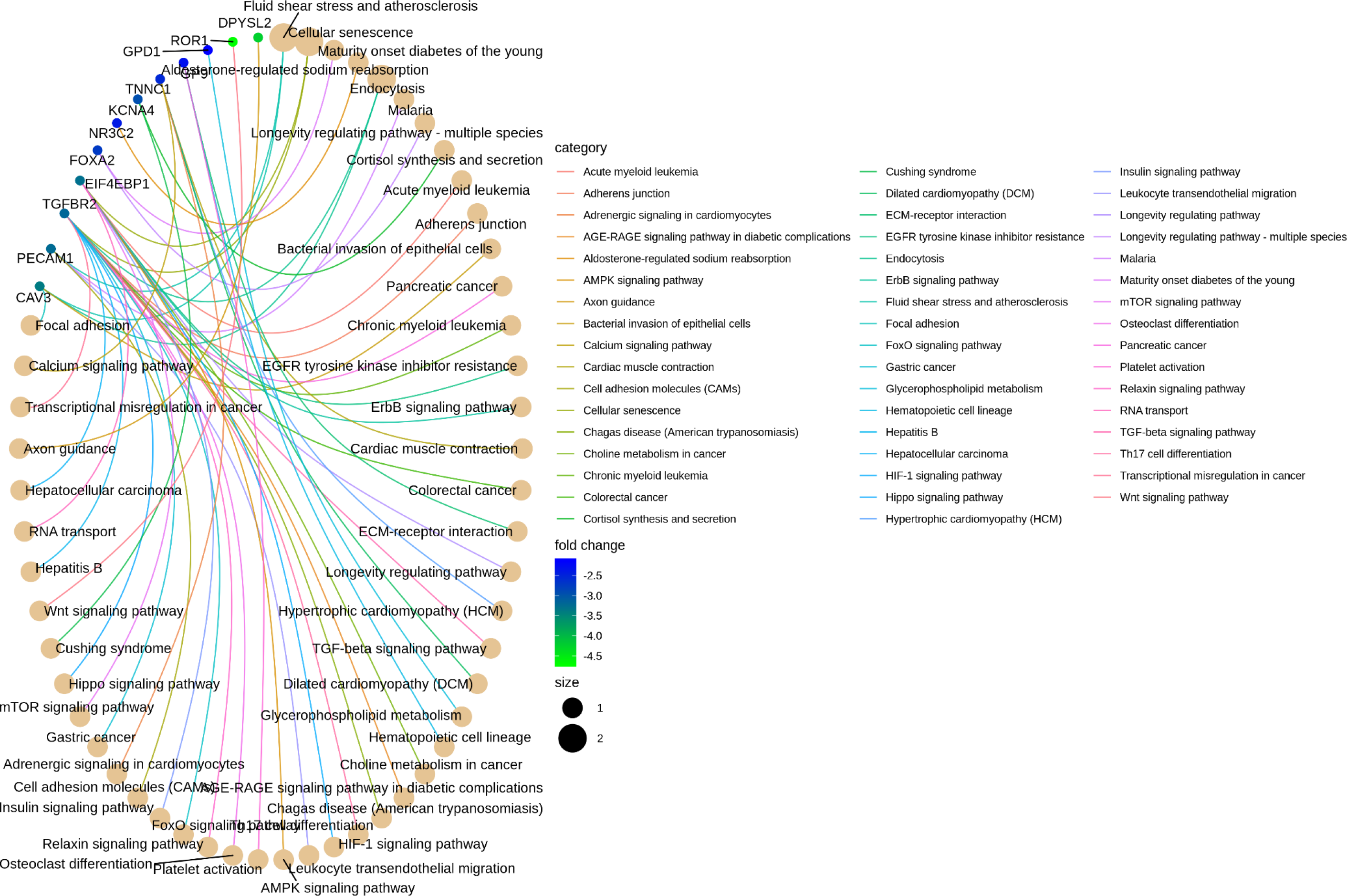
cancer regulatory pathways in Subtype-1 samples

**Figure 10:**
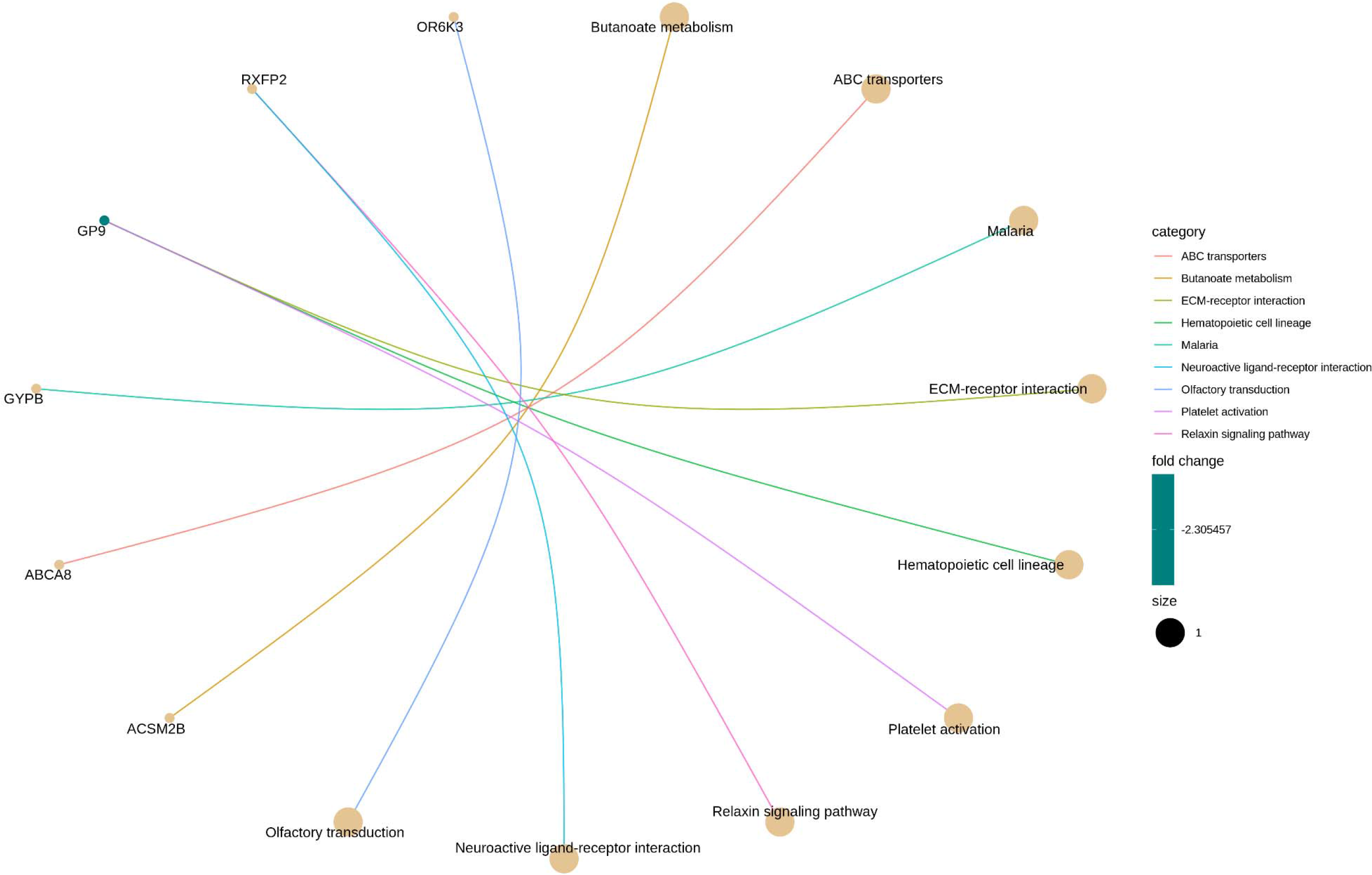
cancer regulatory pathways in Subtype-2 samples

### Validation with survival analysis

The LUSC Tumor samples were classified into two subtypes based on the consensus clustering method. Using the survival data of these samples, survival analysis for the most predictive genes identified by our model in Subtype-1 and Subtype-2 groups was conducted. The genes predicted by our LASSO model accurately predicted the outcome of a patient’s survival using gene expression data. Genes such as GAL, TFAP2A, AFF3, TNNC1, TGBR2, HELT, and SFTA1P yielded accurate predictions for the risk of LUSC cancer and can be used in cancer prediction. Survival plots and its p-value is shown in *(Fig 11 and 12, Supplemental Figure S9)*.

**Figure 11:**
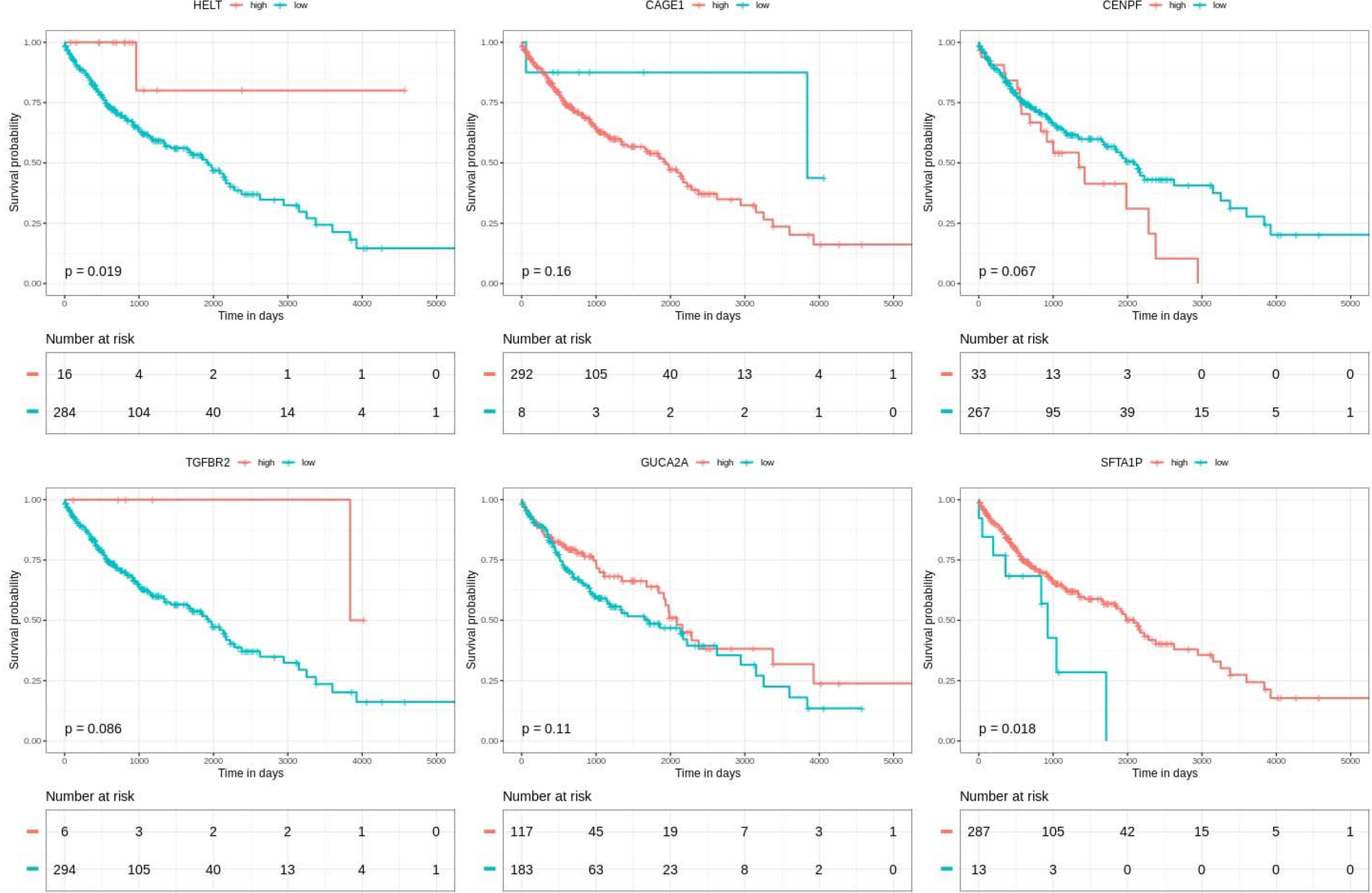
Survival analysis for LASSO predicted genes in Subtype-1 samples

**Figure 12:**
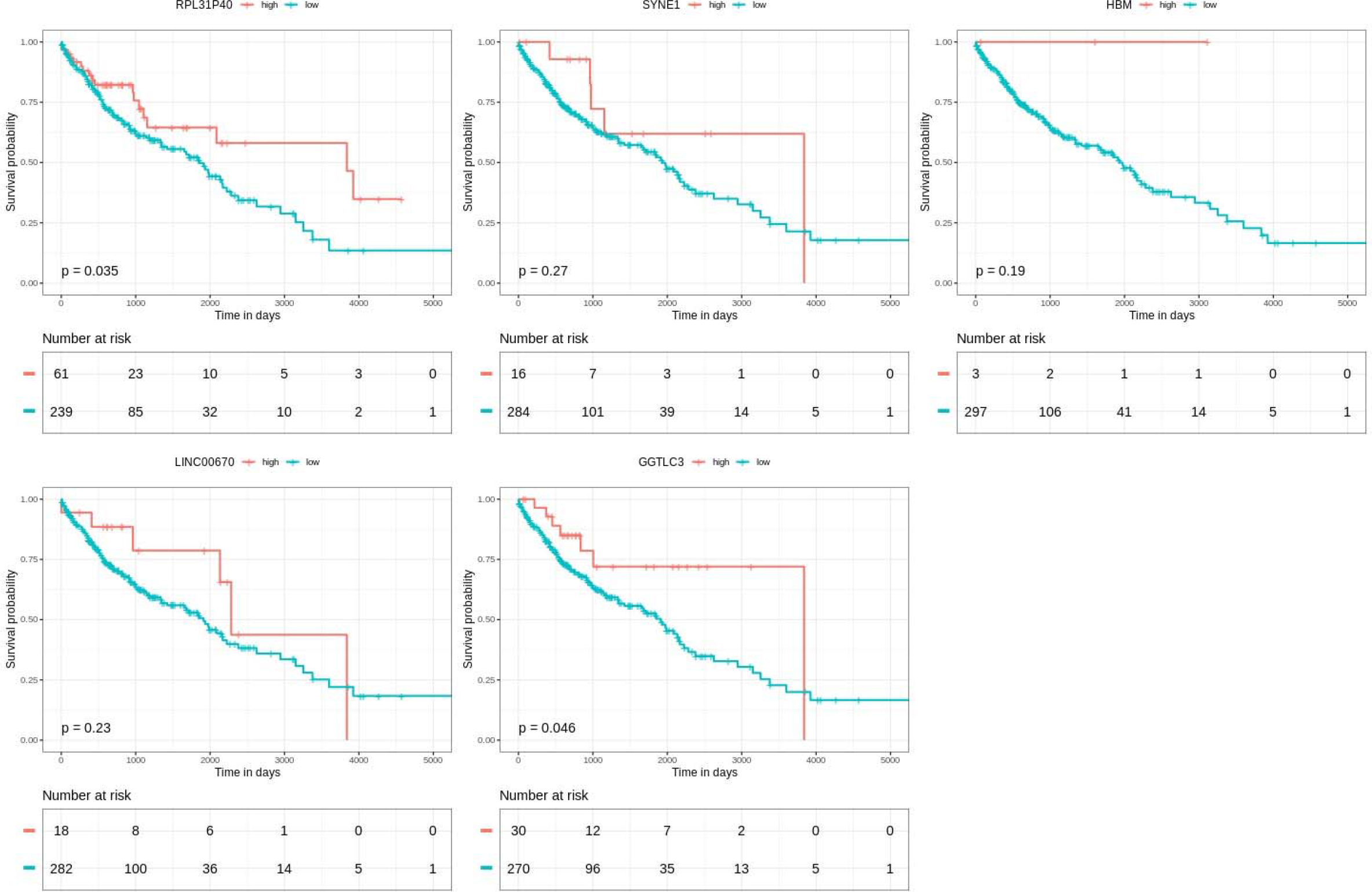
Survival analysis for LASSO predicted genes in Subtype-2 samples

## Discussion

In this study, we developed a LASSO based model for accurate feature selection in LUSC cancer. Our model removed variables that are redundant and removed features which do not add any valuable information in disease prediction. Analysis using the survival data for the predicted genes showed that the model could effectively predict genes responsible for disease prognosis in high dimensional datasets. Deciphering cancer heterogeneity is very critical in understanding cancer dynamics and also for the development of personalized cancer treatment (R. Rosenthal, N. McGranahan, J. Herrero, 2017; Dagogo-Jack & Shaw, 2018) We used Consensus clustering method to determine the number of clusters in our samples, and we clustered the samples into two groups which produced optimal silhouette width for the determined clusters. Differential gene expression analysis showed distinct expression patterns in Subtype-1 and Subtype-2. The number of differentially expressed genes were very high and in these situations, it is difficult to predict the relevant variables. LASSO model was built around 6081 and 8240 DE genes in Subtype-1 and Subtype-2 respectively. Not all the expressed genes were relevant, our model predicted the most relevant genes which were involved in disease progression. Decreased expression of AFF3, TNNC1, TGFBR2, FFAR4 and HELT in Subtype-1 and GGTLC3, GUCA2A, HBM, SFTA1P and SYNE1 in Subtype-2 showed worse overall survival in LUSC cancer samples. Whereas increased expression of genes such as PPIAP45, CAGE1, TFAP2A, CENPF in Subtype-1 decreased overall survival in LUSC cancer samples.

Long intervening noncoding RNAs (lncRNAs) are known to be key regulators of numerous biological processes, and substantial evidence supports that lncRNA expression plays a significant role in tumorigenesis and tumor progression (Ming-Chun Jiang,Jiao-Jiao Ni, Wen-Yu Cui,1 Bo-Ya Wang, 2019). Increased expression of LINC01977, LINC01572 in Subtype-1 samples correlates with worse survival in LUSC cancer subtypes. Whereas, decreased expression of LINC02058 in Subtype-1 and LINC00670 in Subtype-2 showed worse survival in LUSC samples. The LASSO method predicts the most relevant and distinct genes from Subtype-1 and Subtype-2 samples which might be important factor in cancer diagnosis.

The best predictors for subtype 1 and subtype 2 from the LASSO model were found to be involved in several regulatory pathways. The genes such as TGFBR2, EIF4EBP1, and ROR1 which are predicted only in case of Subtype-1 are found to be involved in several cell cycle and growth regulatory pathways and thereby having a strong correlation with cancer. The gene gp9 plays an important role in ECM-receptor interactions, which is critical in disease progression and malignant cell behavior (Walker, Mojares & Del Río Hernández, 2018). The neuroactive ligand-receptor interaction signaling pathway is a collection of receptors and ligands on the plasma membrane that are associated with intracellular and extracellular signaling pathways. It is found to be associated with prostate cancer, bladder cancer, and renal cell carcinoma (He et al., 2018). In our study, the gene RXFP2 that is predicted only in Subtype-2 is found to be involved in neuroactive ligand-receptor interaction. RXFP2 is also found to be involved in Relaxin signaling which induces cell invasion and is reported in several cancers (Fue et al., 2018). The modulators of ABC transporters is reported to have the potential to augment the efficacy of anticancer drugs (Chen et al., 2012). ABCA8 is one such gene and it was predicted only in Subtype-2.

Our model identified cancer/testis antigen gene CAGE-1 which is overexpressed in Subtype-1 and might act as a plasma biomarker for lung cancer early detection. Previous studies showed that CAGE-1 provides an important addition to the armamentarium the clinician to aid early detection of lung cancer in high-risk individuals (Park et al., 2003; Kunze, Wendt & Schlott, 2006; Parmigiani et al., 2006; Kunze & Schlott, 2007; Chapman et al., 2011; Kim et al., 2013). GUCA2A was down-regulated in Subtype-2 samples, many studies on GUCA2A indicates its role as a biomarker in diagnosing cancer. Aberrantly expressed GUCA2A can be a candidate marker of poor prognosis in patients with LUSC and Colorectal cancers, which may be a therapeutic target for precision medicine (Kulaksiz H, Rehberg E, Stremmel W, 2002; Chen YB, Zhu YP, Feng HY, Liu Y, Qian J, Fan YT, 2009; Hui Zhang, Yuanyuan Du, Zhuo Wang, Rui Lou, Jianzhong Wu, 2019) Under expression of TGFBR2 in Subtype-1 samples is associated with poor prognosis, and TGFBR2 is also associated with poor prognosis in cervical cancer (Yang et al., 2017; Yokouchi et al., 2017). CENP-F, a cell cycle-regulated centromere protein, has been shown to affect numerous tumorigenic processes, increased expression of CENP-F in subtype-1 correlates with poor survival. Previous studies demonstrate that CENP-F may serve as a valuable molecular marker for predicting the prognosis of esophageal squamous cell carcinoma patients and nasopharyngeal carcinoma progression(J.-Y. et al., 2010; Chen et al., 2011; Mi et al., 2013; Yang et al., 2017). Downregulation of SFTA1P in Subtype-2 correlates with poor survival, previous studies suggest SFTA1P regulates both oncogene and tumor suppressor genes during the carcinogenesis of lung squamous cell carcinoma(Zhao, Luo & Jiao, 2014; Huang et al., 2017; Zhang et al., 2017; Ma et al., 2018) which can be used as a prognostic biomarker.

Furthermore, Consensus clustering and LASSO helps us to choose a model with the most relevant features. Consistent with this finding, the clustered samples into two different subtypes showed distinct features, highlighting the better sample grouping and risk assessment. Moreover, the results of survival analysis validates that the survival time of the predicted genes correlates with gene expression pattern, which is recognizably different in both the Subtypes, indicating that this model could effectively distinguish the samples with different expression pattern by overcoming the feature selection problem and was accurate for predicting the risk of LUSC cancer.

## Conclusions

In conclusion, this study suggests that the unsupervised method such as Consensus clustering and LASSO model-based feature selection could be used to evaluate prediction and prognosis of LUSC cancer. With this model, we can identify the prognostic biomarkers of LUSC cancer, and the model-predicted genes would be helpful for clinicians in the management of cancer patients.

## Acknowledgements

We wish to acknowledge Sangram Keshari Sahu from Bioinformatics lab of the IISER Mohali, India for his respective technical and scientific expertise. The efforts of Sangram in Gene Expression analysis and the expression project for Oncology are greatly acknowledged.

## Notes

https://bitbucket.org/lusc_data/supporting_data/src/master/

