## Supplementary figures and images for "Gene prediction in heterogeneous cancer tissues and establishment of Least Absolute Shrinking and Selection Operator model of lung squamous cell carcinoma"

### Fig S1

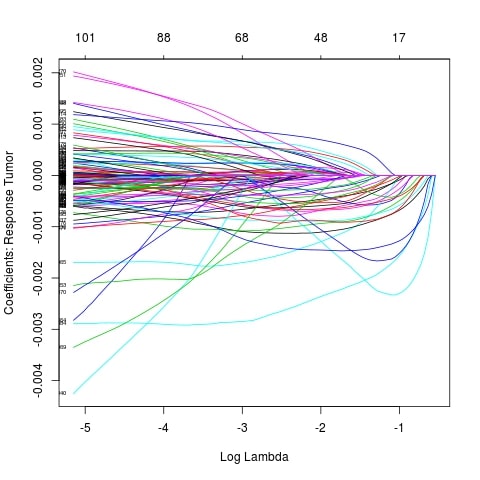

### Fig S3

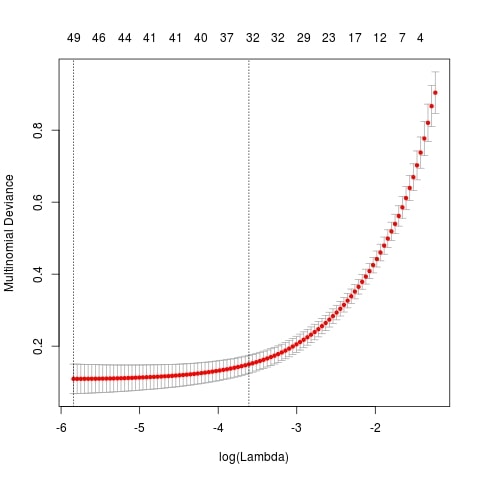

### Fig. S5

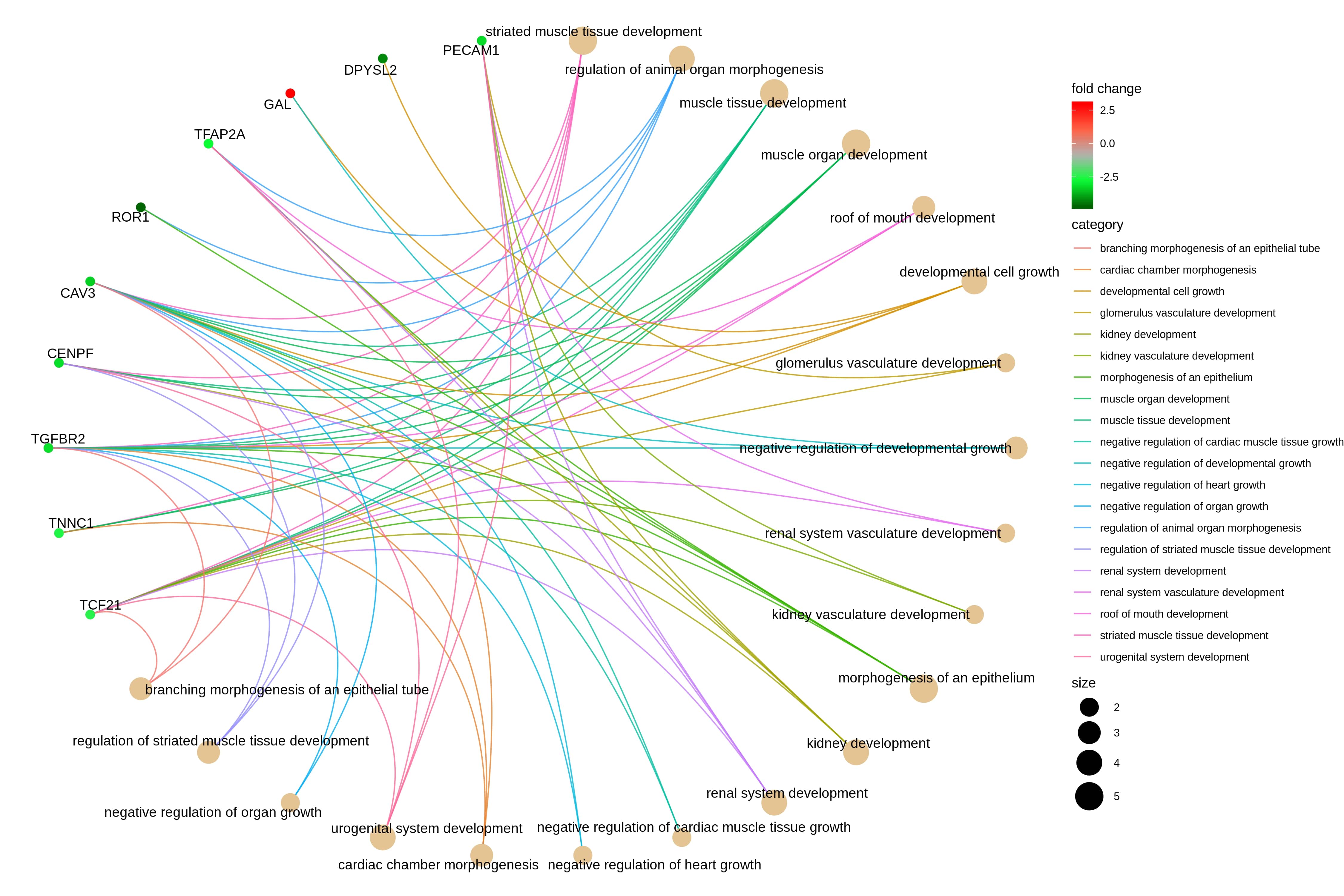

### S2

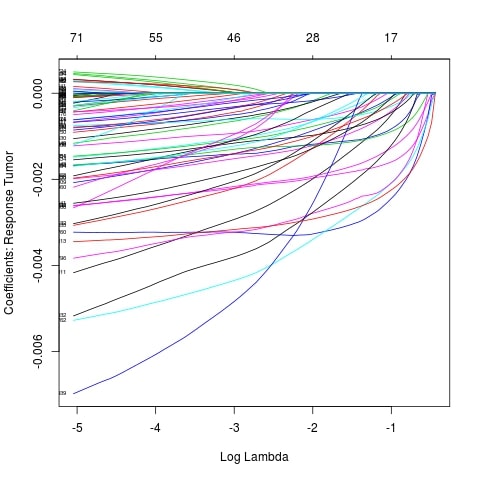

### S4

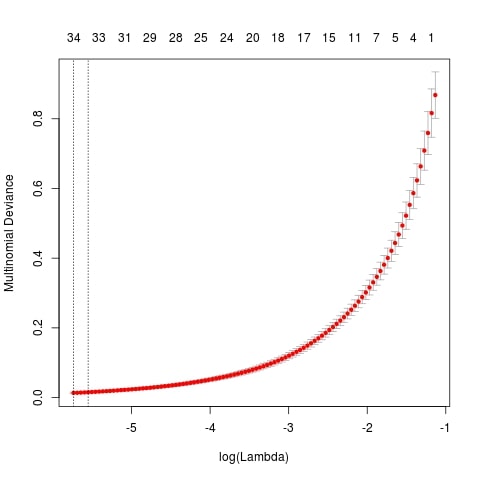

### S6

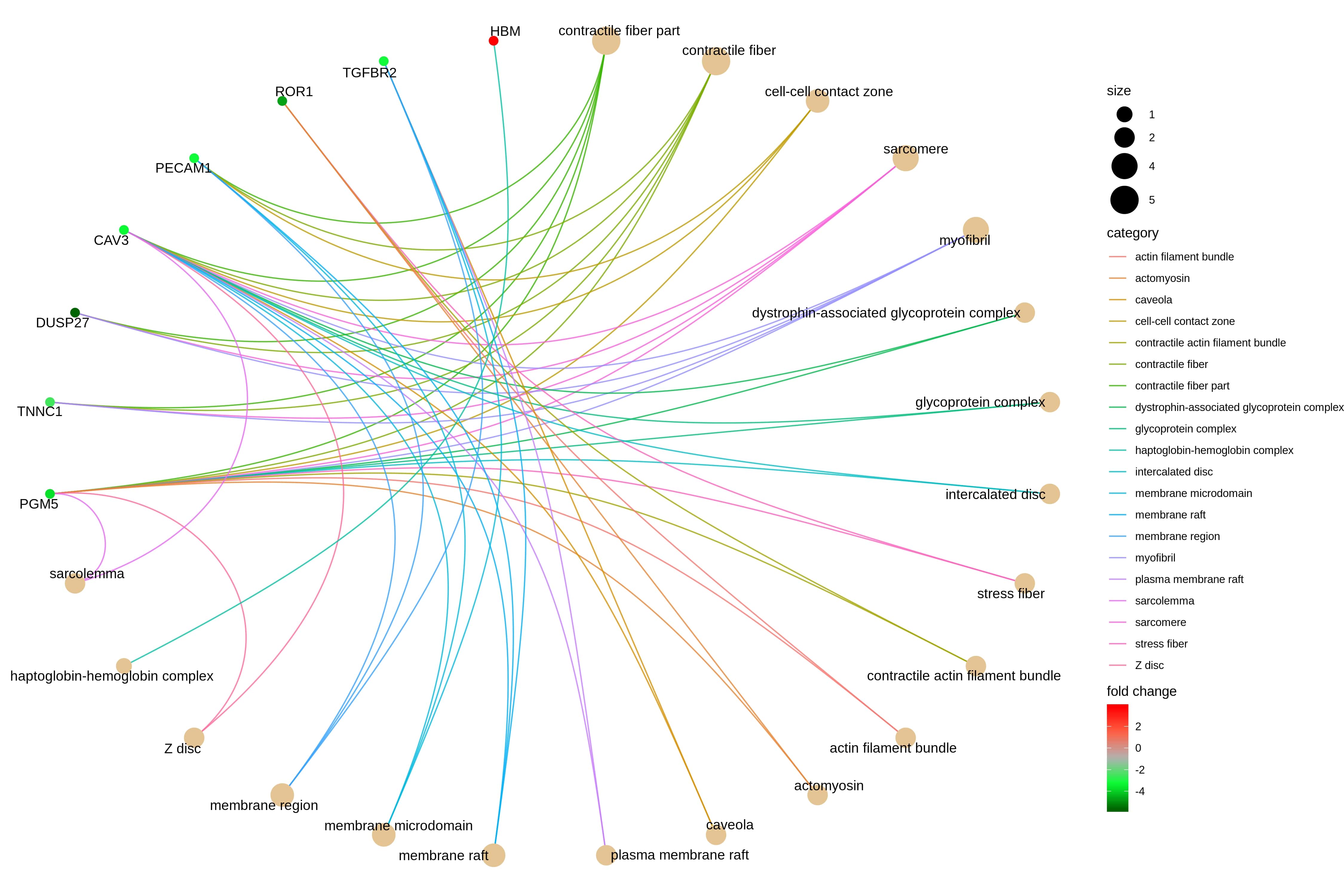

### S7

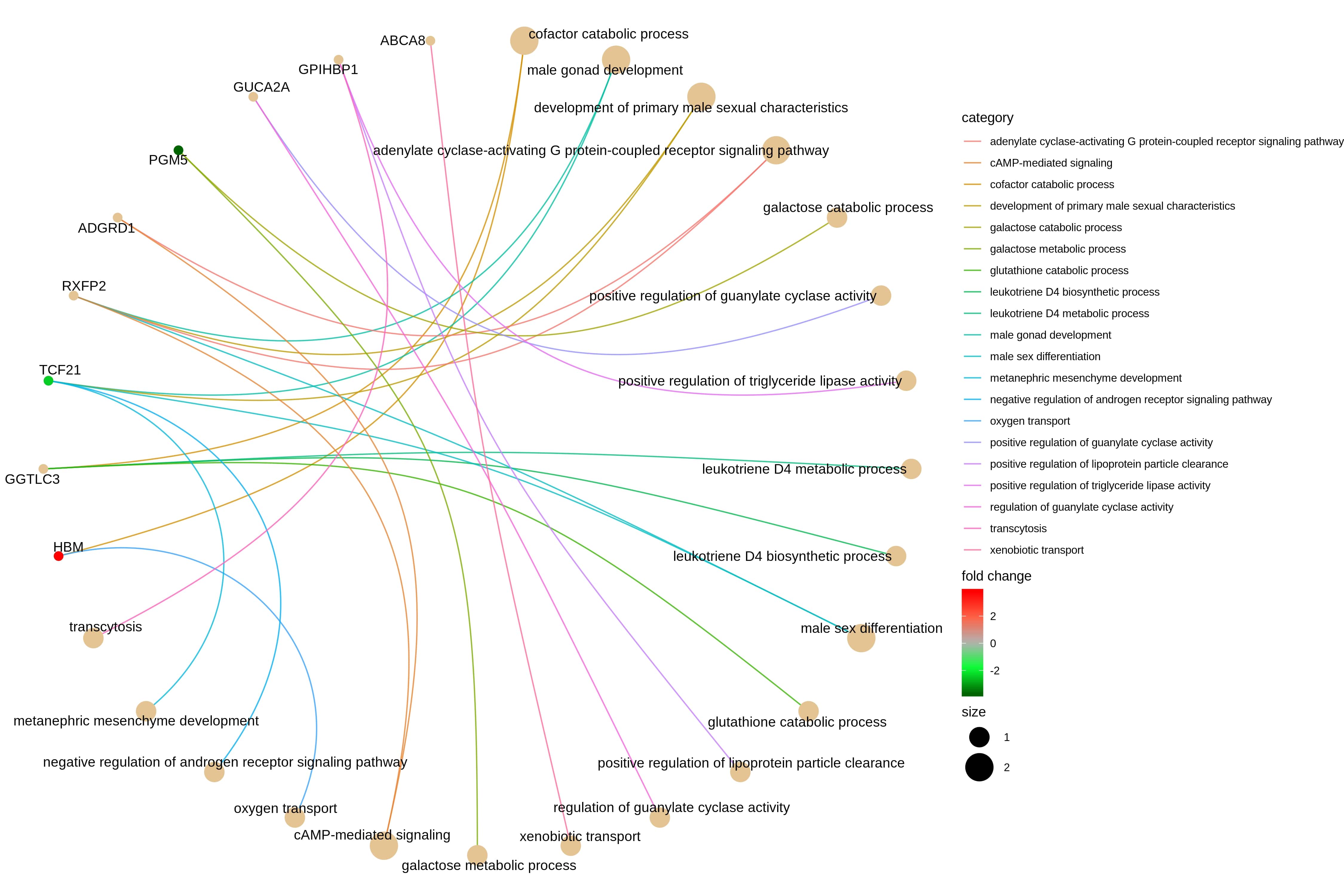

### S8

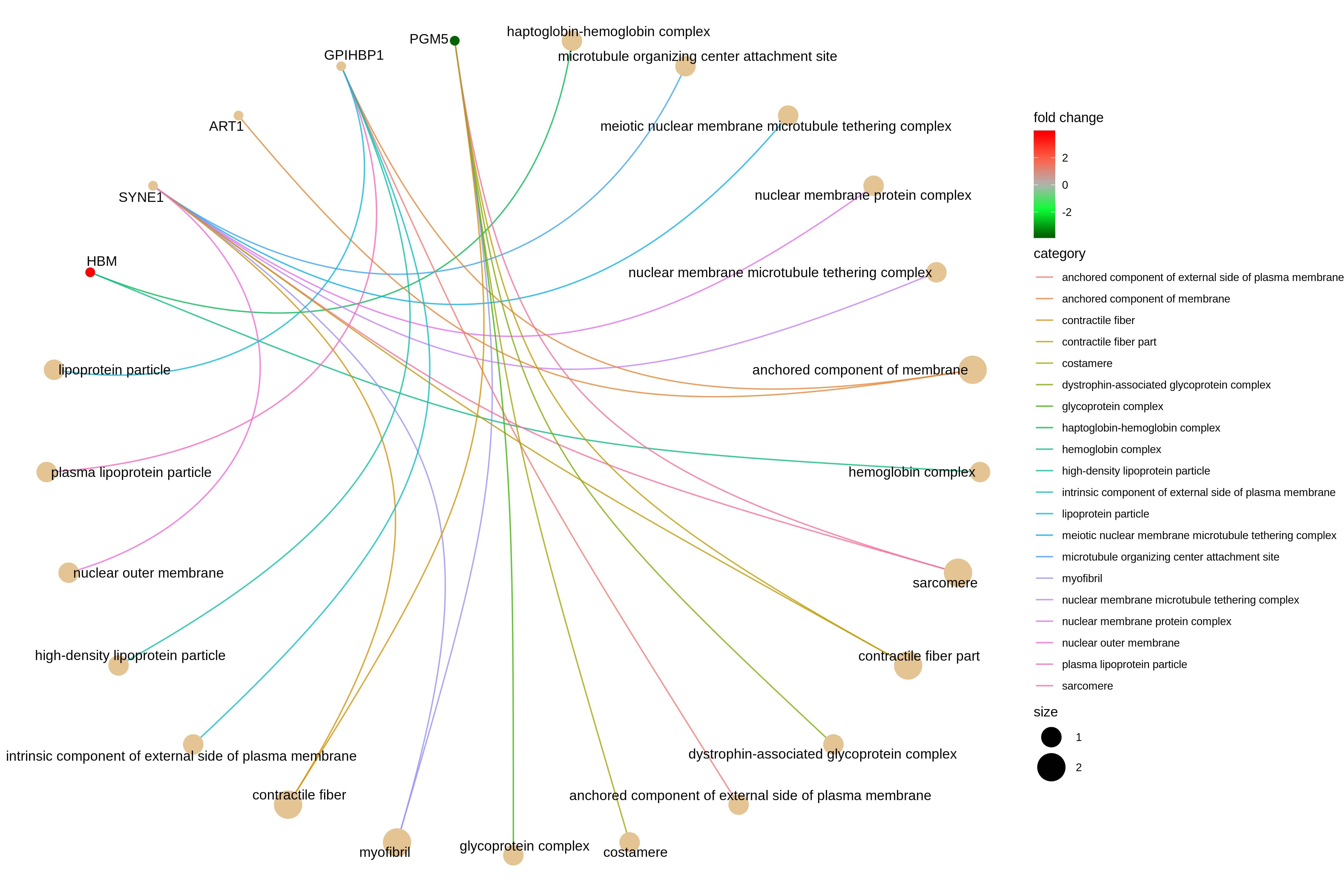
